# Single-cell Multiomics Reveals Clonal T-cell Expansions and Exhaustion in Blastic Plasmacytoid Dendritic Cell Neoplasm

**DOI:** 10.1101/2021.12.01.470599

**Authors:** Erica A. K. DePasquale, Daniel Ssozi, Marina Ainciburu, Jonathan Good, Jenny Noel, Martin Villanueva, Charles P. Couturier, Alex K. Shalek, Sary F. Aranki, Hari R. Mallidi, Gabriel K. Griffin, Andrew A. Lane, Peter van Galen

## Abstract

The immune system represents a major barrier to cancer progression, driving the evolution of immunoregulatory interactions between malignant cells and T-cells in the tumor environment. Blastic plasmacytoid dendritic cell neoplasms (BPDCN), a rare acute leukemia with plasmacytoid dendritic cell (pDC) differentiation, provides a unique opportunity to study these interactions. pDCs are key producers of interferon alpha (IFNA) that play an important role in T-cell activation at the interface between the innate and adaptive immune system. To assess how uncontrolled proliferation of malignant BPDCN cells affects the tumor environment, we catalog immune cell heterogeneity in the bone marrow (BM) of five healthy controls and five BPDCN patients by analyzing 52,803 single-cell transcriptomes, including 18,779 T-cells. We test computational techniques for robust cell type classification and find that T-cells in BPDCN patients consistently upregulate interferon alpha (IFNA) response and downregulate tumor necrosis factor alpha (TNFA) pathways. Integrating transcriptional data with T-cell receptor sequencing via shared barcodes reveals significant T-cell exhaustion in BPDCN that is positively correlated with T-cell clonotype expansion. By highlighting new mechanisms of T-cell exhaustion and immune evasion in BPDCN, our results demonstrate the value of single-cell multiomics to understand immune cell interactions in the tumor environment.

## 1 Introduction

Innovations in immuno-oncology, such as immune checkpoint blockade (ICB) therapy, have transformed cancer medicine. ICB and CAR-T cells have led to improved outcomes in various solid tumors and B-cell malignancies, respectively (1), however these methods have been less successful in myeloid leukemias than in other cancers. Various mechanisms have been proposed to explain the relative inefficacy of immunotherapies in myeloid leukemias, including expression of immunoregulatory molecules by malignant cells (2,3). The complexity of immune regulation in cancer is well illustrated by expression of interferon (IFN) related genes that influence the effectiveness of immunotherapy through immunostimulatory and immunosuppressive effects (4–6). Chronic interferon signaling has been associated with T-cell exhaustion in the setting of viral infections, but its role in cancer is controversial (7–9). Blastic plasmacytoid dendritic cell neoplasm (BPDCN) is an aggressive form of acute leukemia with few effective therapies that provides a unique opportunity to study IFN dysregulation in cancer. BPDCN is characterized by uncontrolled proliferation of transformed plasmacytoid dendritic cells (pDCs), specialized immune cells that link the innate and adaptive immune systems through the secretion of Type I interferons, including IFNA, particularly during viral infection (10). Studies of BPDCN up to this point have largely focused on the malignant pDC-like tumor cells, but few have focused on the T-cell response. An effective immune response relies on the interaction between healthy innate and adaptive immune systems, which is critical for immunotherapy and may be impacted by IFNA-producing pDCs.

Single-cell RNA-sequencing (scRNA-seq) has provided granular insights into the dynamics and phenotypes of immune cells. Specifically, scRNA-seq has been utilized to define subsets of exhausted T-cells in viral infection and cancer, including those that drive responses to ICB (11–13). Advances in other single-cell technologies — such as single-cell methylation, chromatin accessibility, and mutation status — have allowed for multimodal data collection all from the same cell (14–17). This type of analysis allows us to understand relationships between biological processes and heterogeneous cell types that cannot be studied using unimodal measurements alone. To study T-cell biology, scRNA-seq can be paired with sequencing of the T-cell receptor (TCR) α- and β-chain variable regions. These regions can be enriched from single-cell transcriptomes while maintaining cell barcodes to integrate TCR sequencing data with gene expression profiles (18–20). Recently, TCR sequencing has been applied to solid and blood cancers to tie cell phenotypes to TCR properties, which has allowed for a deeper examination of T-cell subsets via multiple modalities (20– 22). These studies also investigate the phenomenon of T-cell exhaustion, which occurs when cells enter a dysfunctional state in which they lose effector functions, such as proliferation and cytotoxicity, and gain immunoregulatory functions (7,23,24). Known drivers of T-cell exhaustion include continuous antigenic stimulation and chronic inflammation (7). Understanding the mechanism by which cells become exhausted is particularly important in the context of immunotherapy, as exhausted T-cells can regain effector functions via ICB therapy (24,25).

Here, we investigate the immune environment in five BPDCN bone marrow samples and five healthy controls using scRNA-seq and TCR sequencing. We used computational integration tools to create a unified healthy reference and applied multiple classification algorithms to annotate cell types in the BPDCN samples by consensus. These data show variable cell type expansions and a wide range of T-cell proportions between patients. Though we found heterogeneity in cell type proportions, we identified common signatures of upregulated IFNA response, downregulated TNFA signaling, and significantly increased CD8+ T-cell exhaustion. We applied a TCR sequencing approach termed T-cell Receptor Enrichment to linK clonotypes (TREK-seq) that we recently developed to identify expanded T-cell clonotypes in BPDCN samples and showed a correlation between CD8+ T-cell clone size and exhaustion scores. By providing a comprehensive map of cellular heterogeneity and T-cell transcriptomes/clonotypes in BPDCN patients, we lay the foundation for future development and evaluation of immunotherapies in this devastating cancer.

## 2 Materials and Methods

### 2.1 Patient samples

Bone marrow aspirate was collected from 5 patients with blastic plasmacytoid dendritic cell neoplasm (BPDCN) who consented to an excess sample banking and sequencing protocol that covered all study procedures and was approved by the Institutional Review Board (IRB) of the Dana-Farber/Harvard Cancer Consortium. Additionally, 5 samples from healthy donors were collected for use as a control in this study, following the same approved protocol or a protocol approved by the IRB of Mass General Brigham. Details for each of these samples are located in **Supplemental Table 1**.

### 2.2 Single-cell RNA-sequencing

Single-cell RNA-sequencing was performed on cryopreserved iliac crest bone marrow aspirates for BPDCN samples and BM 1-3 controls, and cryopreserved sternum bone marrow for BM 4 and 5 controls. To isolate mononuclear cells, BPDCN samples and BM 1-4 were processed using Ficoll or lymphoprep, whereas BM 5 was processed using Acrodiscs (Pall AP-4952). Cells were stored in liquid nitrogen, thawed using standard procedures, and viable (DAPI negative) cells were sorted on a Sony SH800 flow cytometer. Next, 10,000-15,000 cells were loaded onto a 10x Genomics chip. Further processing was done using the recommended procedures for the 10x Genomics 3’ v3.0 or v3.1 chemistry (26). 10x libraries were sequenced on the NovaSeq SP 100 cycle with the following parameters (Read 1: 28 + Read 2: 75 + Index 1 (i7): 10 + Index 2 (i7): 10).

### 2.3 Dataset processing

Raw sequencing data were processed using CellRanger software (27) to generate FASTQ files and count matrices. Ambient RNAs were estimated and removed from the datasets using SoupX with default parameters (28). Each dataset was filtered to retain cells with >= 500 UMIs, >=1000 genes expressed, and <30% of the reads mapping to the mitochondrial genome. Three of the healthy control bone marrow samples were originally integrated using the IntegrateData() function in Seurat v4.0.3 and clustered using the same software at the default resolution of 0.5 (29). Cell cycle genes (“cc.genes” within the Seurat software) were removed from the integration anchors to combine smaller cell cycle-driven clusters and to eliminate redundancy, though these genes were retained in the final dataset for further analyses. Further, a combination of high resolution sub-clustering with Seurat and analysis of key T-cell gene expression were used to improve granularity in the naive T-cell compartment. Two additional healthy samples from older individuals were further integrated into the original controls using Seurat’s TransferData() function. The BPDCN samples were processed and clustered individually using Louvain clustering within Seurat with default options and a resolution of 0.5, though these clusterings were not used in downstream analyses.

### 2.4 Cell classification

Cells in each BPDCN sample were classified with four classification methods using the integrated healthy control samples as a reference: random forest (30,31), cellHarmony (32), Seurat’s TransferData() function (33), and scPred with default parameters (34). Reference input for three of the algorithms was the integrated Seurat object and feature selection was performed separately as a part of each method. cellHarmony instead used a gene expression matrix of the cells by top 50 marker genes for each cluster (as defined by Seurat), the full expression matrix, and a table to cell type classifications for the reference. Given highly consistent classifications between the four methods, we selected cellHarmony-defined cell type labels for all downstream analyses. Classified cells from BPDCN patients were projected into the UMAP space of the integrated controls using the MapQuery() function in Seurat v4.0.3.

### 2.5 Gene Set Enrichment Analysis (GSEA)

Gene set enrichment was performed separately on cells from each cell type and BPDCN sample relative to the healthy controls. Log fold change values for every gene were extracted from the cellHarmony output (∼/input/cellHarmony/OtherFiles), sorted in decreasing order, and used as input for a custom GSEA function (35). This function uses two GSEA implementations in R, gage (36) and fgsea (37) and only reports the results that are significantly enriched with both methods. Hallmark gene sets from the GSEA website were used for this analysis (https://www.gsea-msigdb.org/gsea/msigdb/collections.jsp). Pathways that were significantly enriched in at least one comparison were plotted using the Pheatmap R package.

### 2.6 Pathway score quantification

To score individual cells for gene signatures, we combined all 52,803 cells from the dataset into one Seurat object, read in a curated list of signatures (IFNA response from the GSEA Hallmark gene set, TNFA signaling by NFKB from the Hallmark gene set, T-cell exhaustion from Penter et al. (21)) and applied the function AddModuleScore to the Seurat object using signature genes as features. Statistical significance for each cell type was calculated by comparing the median scores of normal samples (n = 5) to the median scores of BPDCN samples (n = 5) using the R function wilcox.test and default parameters. The number of biological replicates was less than five if a cell type was not detected in one of the donors (for example, n = 4 for BPDCN CD8+ Memory T-cells).

### 2.7 T-cell receptor sequencing

T-cell receptor sequencing was performed on both control and BPDCN bone marrow samples using a modified protocol originally developed for Seq-Well (18). This protocol adaptation termed TREK-seq is described in an independent manuscript (Miller et al. Nat Biotechnol., under review). The modifications to the original protocol are as follows: in the TCR enrichment master mix, we added PartialRead1 and PartialTSO primers at a final concentration of 1.25 µM each. For amplification of TCR transcripts following enrichment, we used the same primers at a final concentration of 0.4 µM each. For the final PCR, we used UPS2-N70x and 10X_SI-PCR_P5 primers at a final concentration of 0.2 µM each to add the Illumina P5 and P7 sequences. The libraries were sequenced using a 150 cycle kit on the Illumina MiSeq and loaded at a final concentration of 10 pM. 28 cycles were used for Read 1, which read the cell barcode and UMI. 150 cycles were used for Index 1, which read the TCR region. TCRα and TCRβ-specific custom sequencing primers were used for index 1 at a final concentration of 2.5 µM. We aimed for a MiSeq cluster density of roughly 450k/mm2. Primer sequences can be found in **Supplemental Table 2**.

### 2.8 TREK-Seq computational pipeline

Raw sequencing data were demultiplexed with bcl2fastq (v2.20.0), and the resulting fastq files were reformatted to join the cell barcode, UMI and corresponding TCR sequence in the same file. For this analysis, we developed and applied the bioinformatics pipeline WARPT, or Workflow for Association of Receptor Pairs from TREK-seq (https://github.com/mainciburu/WARPT). Briefly, we first corrected cellular barcodes allowing one mismatch with the 10x Single Cell 3’ v3 whitelist. UMI correction was also performed by clustering together UMIs with one mismatch. Every corrected barcode and UMI sequence was then added to the corresponding fastq read header. Next, we applied a quality filter to remove every read with an average score < 25. To account for barcode swapping, TCR sequences with identical barcode and UMI were subjected to clustering using an identity threshold of 0.9. The subsequent analysis was carried out exclusively with clusters representing at least 50% of the reads with identical barcodes and UMI and doubling the number of reads from the second most abundant cluster. Next, a consensus sequence was built to summarize the TCR sequences in each of those clusters. We required a minimum of 3 sequences and allowed for a maximum error rate of 0.5 and gap frequency of 0.5 per position. Consensus sequences were aligned to the VDJ segments reference available at the IMGT database. IgBLAST with the default parameters was used for the alignment. Finally, the V, D and J calls and CDR3 sequence with higher UMI counts, for both *TRA* and *TRB* genes, were assigned to each cell barcode. For downstream analysis, only *TRB* variable regions were used.

### 2.9 Statistics

P-values less than or equal to 0.05 were considered statistically significant. Cell population proportions between younger and older healthy donors were assessed for significant differences using a chi-squared test. For the gene set heatmaps, P-values were calculated via t-test for each comparison of reference to BPDCN sample per cell type and adjusted using the Bonferroni method. Genes that were significant following correction and had an expression value of 0.5 or greater (log normalized expression) in at least 1 sample were retained for the heatmap. A Wilcoxon signed-rank test was used for pathway score quantification, a Kruskal-Wallis rank sum test was used to compare normalized T-cell clonotype sizes, and a Spearman ranked correlation analysis was used to test the relationship between clonotype size and exhaustion scores.

### 2.10 Data and software availability

All data used in this paper will be made publicly available in the Gene Expression Omnibus (GEO) (https://www.ncbi.nlm.nih.gov/geo/). The scripts used to generate the results in this paper can be found at the project GitHub repository: https://github.com/EDePasquale/BPDCN.

## 3 Results

### 3.1 Identification of Cell Populations in Healthy Bone Marrow

To map the cellular diversity in healthy bone marrow samples, we performed scRNA-seq using the 10x Genomics platform (27). We profiled cells from three younger healthy donors (age 31-45) and two older donors (age 74 and 75) (**Supplemental Table 1**). The two older donors were included as age-appropriate controls for BPDCN, which has been reported to have a median age of ∼65 (38). Unique cellular barcodes were used to assign transcripts to cells and individual mRNA molecules were quantified using Unique Molecular Identifiers (UMI). Across all five healthy donors, we retained 25,726 cells following quality control and filtering.

Cells from these healthy donors (BM 1-5) were integrated using the Seurat package (IntegrateData and TransferData functions) to remove batch effects and generate a unified UMAP projection of the data (**Figure 1A and Supplemental Figure 1**). We initially identified 16 unique clusters based on gene expression that were interrogated for expression of canonical marker genes and gene signatures of blood cell types (**Figure 1B**). Additional sub clustering was performed in the lymphocytic compartment, and new T-cell classifications were made based on expression of *CD8A, CD4, ITGB1*, and *CCR7* gene expression. Specifically, CD4+ T-cells and CD8+ T-cells could both be split into Naive (*ITGB1-CCR7+*) and Memory (*ITGB1+ CCR7-/+*) subsets. The final reference was comprised of 17 clusters representing known hematopoietic cell types, including HSCs and progenitors, myeloid, erythroid, and lymphoid cells (**Figure 1C**). All cell types were represented in each donor at broadly comparable levels, though variation exists as expected (**Figure 1D**) (39). We observed no statistically significant differences in population proportions between the donors as an effect of age or collection procedure (*P* = 0.2351).

**Figure 1:**
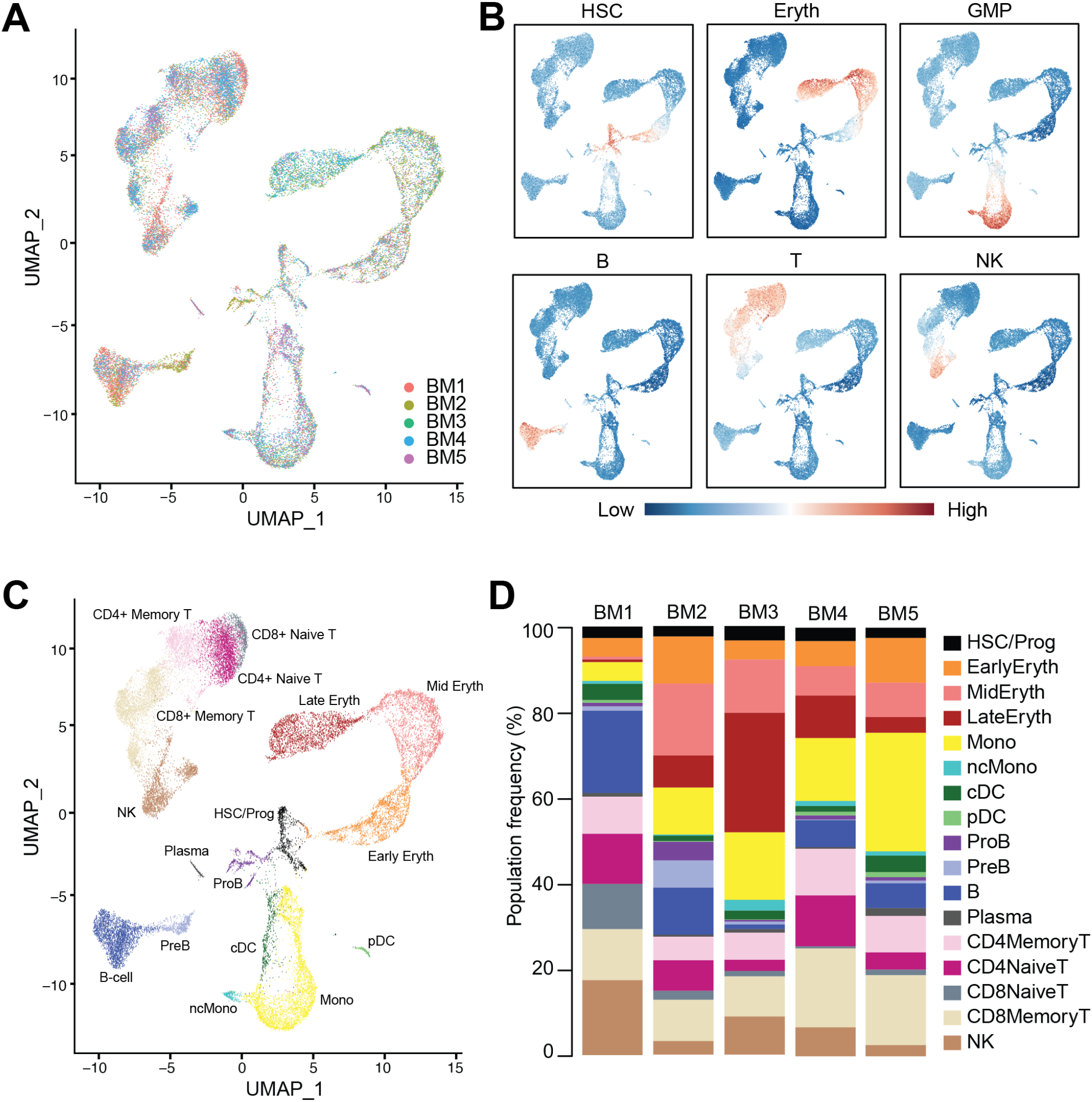
Integration and labeling of five healthy bone marrow control samples. (A) UMAP visualization of Seurat-integrated scRNA-seq data for 25,726 hematopoietic cells from five normal BM aspirates. (B) Expression scores for lineage signatures overlaid on the UMAP of healthy BM in (A). (C) UMAP shows 17 clusters of cells with similar transcriptional states, identified by Seurat clustering and sub-clustering of healthy BM. (D) Stacked barplots show the frequencies of cell types in five normal BMs.

### 3.2 Single-Cell Profiling of BPDCN Bone Marrow

Approximately half of BPDCN patients present with skin tumors only, whereas the other half present with bone marrow involvement, i.e. detection of bone marrow tumor blasts by conventional clinical assays such as morphologic examination or flow cytometry (40). To understand the proportional and transcriptional changes that occur in the bone marrow in the presence of BPDCN tumor cells, we selected five patients with 1.5-97% disease involvement at diagnosis (**Supplemental Table 1**). We performed scRNA-seq on BPDCN bone marrow samples using the same protocol as in the healthy samples. From this sequencing, we acquired 27,077 cells across all BPDCN donors following quality control and filtering.

We next wanted to classify cell types in the five BPDCN samples (BPDCN 1-5) using the healthy controls cells as a reference. To this end, we tested four computational classification methods that utilize different underlying algorithms to assign cell type labels to cells: random forest, cellHarmony, Seurat TransferData, and scPred. The resultant cell type predictions were concordant across all four methods in most cell types, particularly in the erythroid and lymphoid lineages, lending confidence to the classification methods (**Figure 2A**). Some variability was noted in the proportion of cells labeled as pDC, cDC, and ProB in BPDCN 4 and 5; we hypothesized that these could be malignant cells that align poorly to any healthy population, which would be consistent with the clinical annotation of high malignant cell content in these bone marrows (**Supplemental Table 1**). To visualize shifts in gene expression programs and differential cell type proportions, we projected the cells from each BPDCN patient onto the same UMAP space as the healthy reference, colored by cellHarmony labels (**Figure 2B**). Indeed, we observed that the pDC and cDC labeled cells in BPDCN 4 and 5 were shifted toward the HSC/Prog and ProB populations, suggesting that the malignant tumor cells in these samples were transcriptionally different from their healthy counterparts. For this study, we were primarily interested in the role of T-cells in response to and as a potential treatment for BPDCN, therefore we next focused on alterations in cells of the lymphoid lineage.

**Figure 2:**
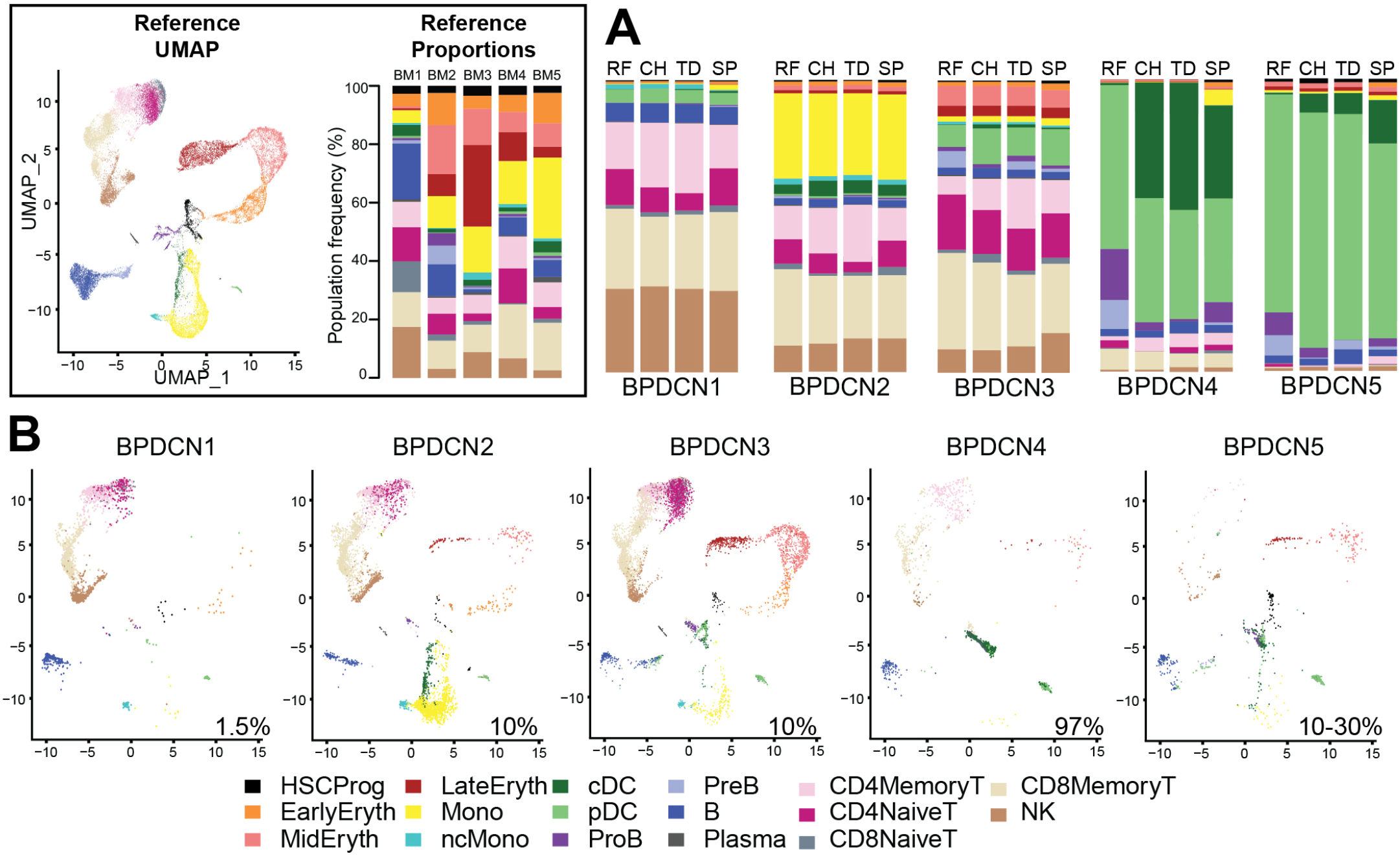
Classification of BPDCN samples using healthy references. (A) Stacked barplots show the frequencies of cell types in five BPDCN samples as classified by four algorithms: RF = random forest, CH = cellHarmony, TD = Seurat TransferData, and SP = scPred. (B) UMAP visualization of cells from five BPDCN samples in the same UMAP space as the integrated reference. Colors in the cell-type proportions and UMAP visualizations are coded by the legend. For comparison with healthy controls, reference UMAP and proportions for each healthy sample are provided in the black box.

### 3.3 T-Cell Populations Show Differential Regulation of IFNA and TNFA Pathways

We used Gene Set Enrichment Analysis (GSEA) to identify pathways that were significantly enriched in the BPDCN samples. Using the Hallmark gene sets and gene expression changes derived via cellHarmony, we generated a heatmap containing all up- or down-regulated pathways enriched in any BPDCN sample compared to the healthy controls (**Figure 3A**). In several T and NK cell populations from BPDCN patients, we found upregulation of “interferon alpha response”, “allograft rejection”, “MYC targets”, and “oxidative phosphorylation”. Interferon alpha (IFNA) response genes were significantly upregulated in 4/4 BPDCN samples in CD8+ Memory T-cells, 3/5 in CD4+ Memory T-cells, 3/3 in CD8+ Naive T-cells, and 3/3 in CD4+ Naive T-cells, along with 3/5 samples in NK cells. IFNA is a marker of immune activation and pDCs produce IFNA, furthering our interest in this pathway. Using a different statistical framework, we next scored all individual CD8+ Memory T-cells for their expression of genes in the IFNA response gene set. We found a significant increase in BPDCN 1-4 relative to control samples, confirming increased IFNA response gene expression in BPDCN T-cells (*P =* 0.0159, **Figure 3B**). BPDCN 5 did not have cells classified as CD8+ Memory T-cells and was not tested. Increased IFNA response gene scores were also observed in CD4+ Memory T-cells and NK cells (data not shown). We found that tumor necrosis factor alpha (TNFA) signaling was downregulated in many of the T-cell and NK populations: 4/4 CD8+ Memory T-cell, 3/5 CD4+ Memory T-cell, and 2/3 CD4+ Naive T-cell, and 3/5 of the NK cell samples with the exception of CD8+ Naive T-cells (0/3) (**Figure 3A**). Scoring individual CD8+ Memory T-cells for the TNFA gene signature also showed significant reductions in BPDCN (*P =* 0.0159, **Figure 3C**). These results demonstrate that T/NK-cells in BPDCN exhibit consistent gene expression changes compared to healthy control cells, including increased response to the pDC-related cytokine IFNA and decreased TNFA signaling.

**Figure 3:**
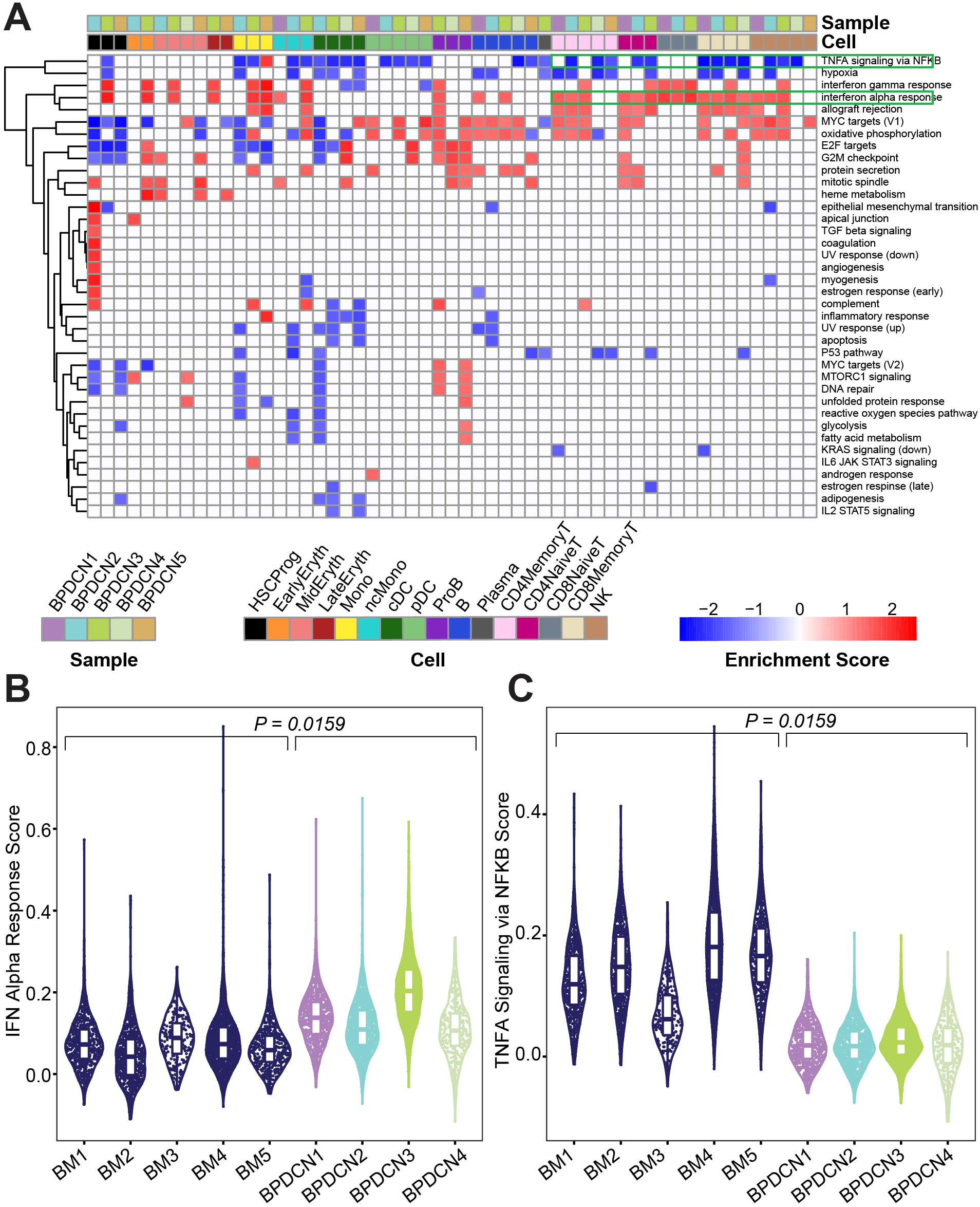
Gene set enrichment shows enrichment of IFNA and depletion of TNFA related genes in BPDCN. (A) Heatmap shows Gene Set Enrichment Analysis enrichment scores for each cell type and sample (columns) and significantly enriched Hallmark pathway (rows), determined by a *P*-value ≤ 0.05 in both GSEA tests. Red indicates high enrichment scores and blue indicates low enrichment scores. (B) Violin plot of IFN Alpha Response gene set scores (y-axis) for CD8+ Memory T-cells from each control and BPDCN sample (x-axis). Each dot within the violin plot represents a cell, with the box in the middle of the violin representing the median and interquartile range of the data. Healthy BM samples are colored in dark blue, BPDCN samples are colored using the sample color scheme from (A). (C) Violin plot of TNFA Signaling via NFKB gene set scores (y-axis) for CD8+ Memory T-cells from each control and BPDCN sample (x-axis). *P*-values were calculated by comparing the medians of n = 5 healthy controls to n = 4 BPDCN samples.

We next investigated which genes in the IFNA response gene set were driving the enrichment in BPDCN T-cells. We generated a heatmap of genes that were significantly different from control expression levels in at least one sample/cell type pair and further filtered to those genes with normalized expression values of at least 0.5 in one pair (p-value ≤ 0.05, Bonferroni adjusted, **Figure 4A**). We also generated heatmaps with the full list of genes in each gene set (**Supplemental Figure 2**). Related to the IFNA response, genes that were upregulated in BPDCN compared to control T-cells in the majority of samples included: *IRF1, PSMB9, PSME2, CD74, ISG15, ISG20, CD47, LY6E, PSMB8*, and *SELL* (**Figure 4A**). The increased expression of *IRF1, PSMB9*, and *PSME2* were notable in all 5 BPDCN samples across lymphoid cell types, with the exception of BPDCN 4’s CD4+ Naive T-cells due to low cell numbers. In contrast, *CD74, ISG15, ISG20, CD47, LY6E, PSMB8*, and *SELL* were most notably increased from control levels in CD8+ Memory T-cells and NK cells, and in BPDCN 1-3 specifically. These genes are involved in T-cell proliferation/differentiation, antigen presentation, and inflammation, consistent with the view that IFNA signaling plays a critical role in immune activation.

**Figure 4:**
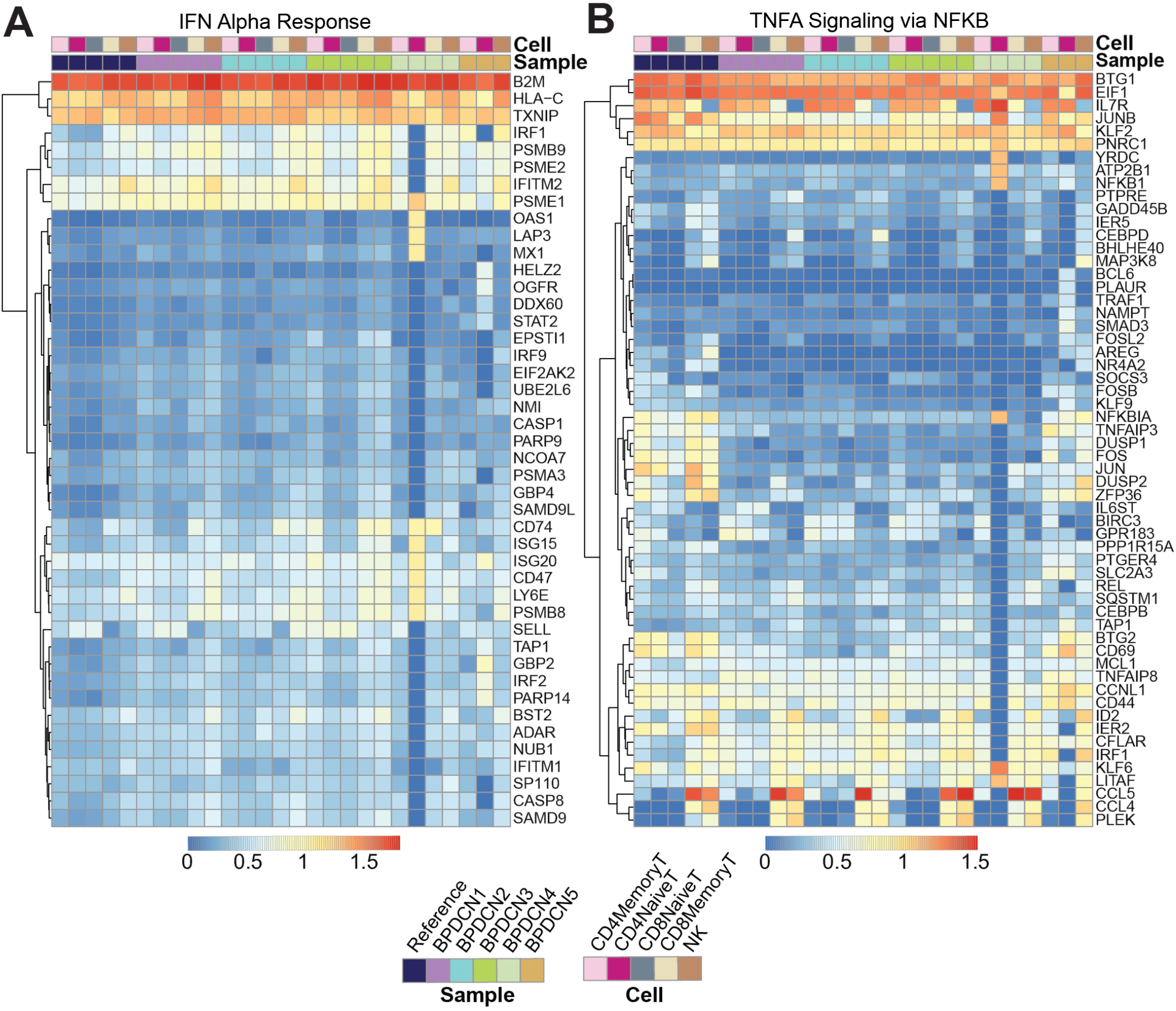
Expression of IFNA and TNFA associated gene sets. (A) Heatmap shows log expression values for genes in the IFN Alpha Response gene set (rows) for each sample and cell type (columns), clustered by row. Genes were filtered to those significant (p < 0.05) after multiple testing correction with an expression value for one sample above 0.5. Red indicates higher expression and blue indicates lower expression. (B) Heatmap shows log expression values for genes in the TNFA Signaling via NFKB gene set.

Related to TNFA signaling, genes that were downregulated in BPDCN T and NK cells compared to control samples are as follows: *NFKBIA, TNFAIP3, DUSP1, DUSP2, FOS, JUN, ZFP36, BTG2*, and *CD69* (**Figure 4B**). These genes were downregulated relative to controls in all T-cell and NK cell subsets in BPDCN 1-4, with the largest expression differences being isolated to the CD4+ and CD8+ Memory T-cell and NK cell clusters. These genes are involved in immune regulation via inflammatory response, proliferation and T-cell activation. Decreased expression of TNFA genes is indicative of a lowered immune response, particularly in relation to T-cell activation. Overall, our results suggest an altered balance between inflammatory pathways in T-cells of BPDCN patients, shifting towards IFNA at the expense of TNFA signaling (41). The changes we observe in IFNA and TNFA target genes are likely to affect the function of the adaptive immune system in BPDCN.

### 3.4 Expanded T-Cell Receptor Clonotypes in BPDCN Correlate with Signatures of Exhaustion

T-cell antigen specificity is determined by the TCR sequence, which consists of the α- and β-chains encoded by *TRAV* and *TRBV* genes that are recombined during T-cell development. We reasoned that sequencing the variable region of the TCR may help to elucidate the biology of T-cells in BPDCN. We performed TCR sequencing on healthy control and BPDCN samples using TREK-seq, a protocol for paired transcript and TCR capture in 10x Genomics data (see **Methods**). We developed and applied the Workflow for Association of Receptor Pairs from TREK-seq (WARPT) computational pipeline to detect groups of cells expressing the same TCR sequence, i.e. T-cell clonotypes. We identified 10,870 T-cell clonotypes in our dataset, each supported by multiple sequencing reads. Next, we ranked clonotypes in each sample by their normalized size (number of cells with the same TCR over all cells in which a TCR was detected). For BM 1-5, we found that none of the control samples were dominated by a single clone or handful of clones, but that the distribution of clonotypes was largely uniform (**Figure 5A-E**). BPDCN samples showed heterogeneity in the distribution of clonotypes: while BPDCN 2-4 do not exhibit a single prevalent clone, similar to the healthy controls, we detected a clone that comprises 30.9% of all TCRs in BPDCN 1 (**Figure 5F-I**). In the healthy controls, TCR sequences were nearly exclusively detected in T-cells (94.7%, **Figure 6A**), and we mainly observed clonal expansions in the CD8+ Memory T-cell compartment, which was expected (**Figure 6B**). These results support that the TREK-seq protocol and analysis pipeline we developed is highly accurate with minimal spurious TCR detection. However, some BPDCN samples showed unexpected cell types containing T-cell receptors (**Figure 6A**). In BPDCN1, one-third of the cells within the largest clone are labeled as NK cells (**Figure 5F**). This sample has a large number of cells classified as NK cells, but a limited tumor cell involvement of 1.5% (**Supplemental Table 1**), leaving open the possibility that the NK-labeled population with TCR results are NKT cells. In BPDCN 4, we unexpectedly captured a minor fraction of rearranged TCR sequences in cDCs (**Figure 5I**). While a rearranged TCR has been detected in tumor cells of some CD4+CD56+ hematologic malignancies (42), this has not been observed in BPDCN, illustrating the potential of single-cell multiomics to reveal unexpected features in the tumor environment.

**Figure 5:**
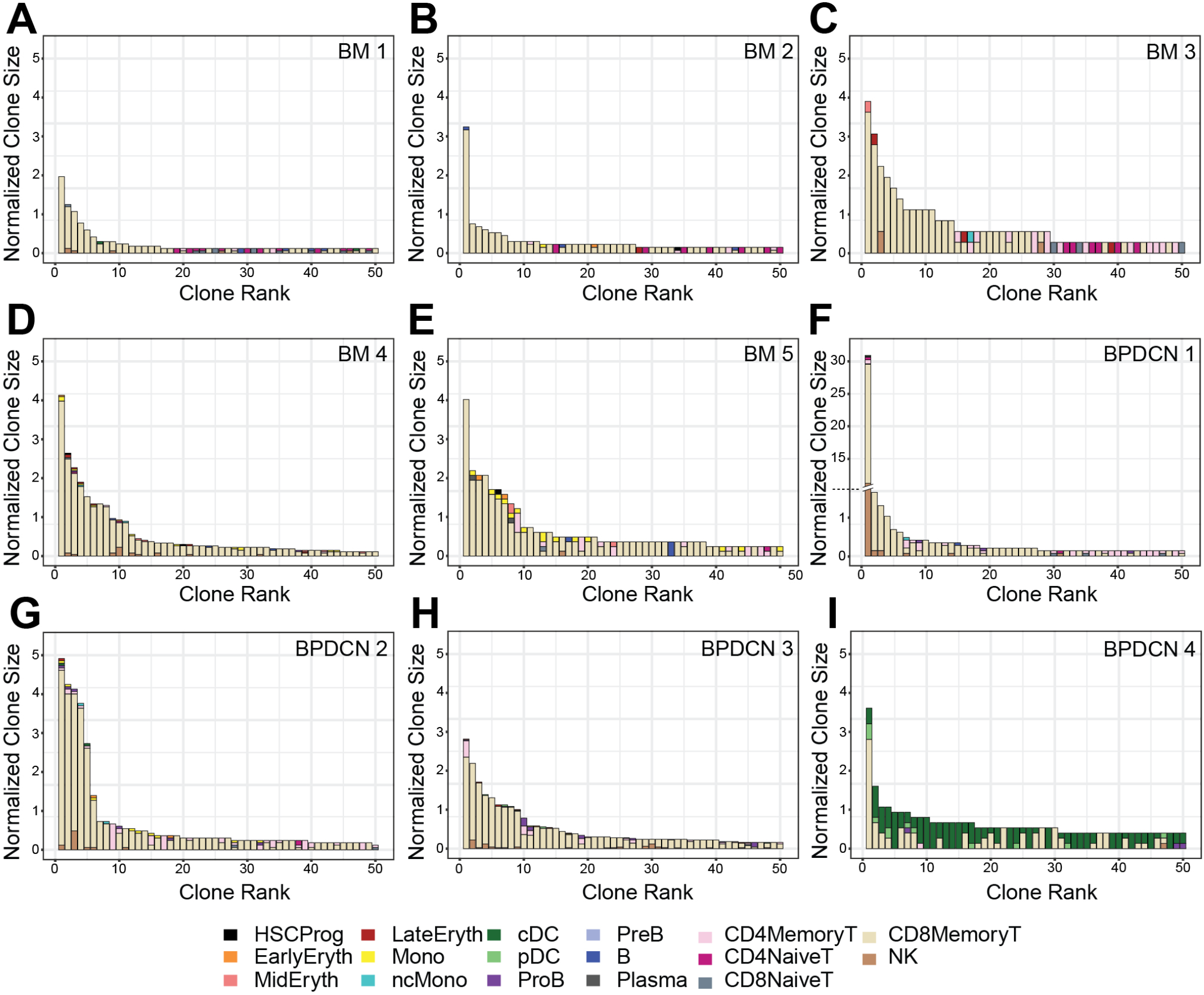
T-cell clonotype sizes in healthy controls and BPDCN patients. (A-E) Bar plots of the top 50 ranked normalized clone sizes for BM 1-5 healthy controls. The normalized clone size is the percentage of cells expressing the same TCR of all cells in which a TCR was detected. For each clone (stacked bar), the colors indicate the proportion comprised of that cell type. (F-I) Bar plots of ranked clones for BPDCN 1-4 patient samples. Axis break in (F) is indicated by dashed line and broken bar.

**Figure 6.**
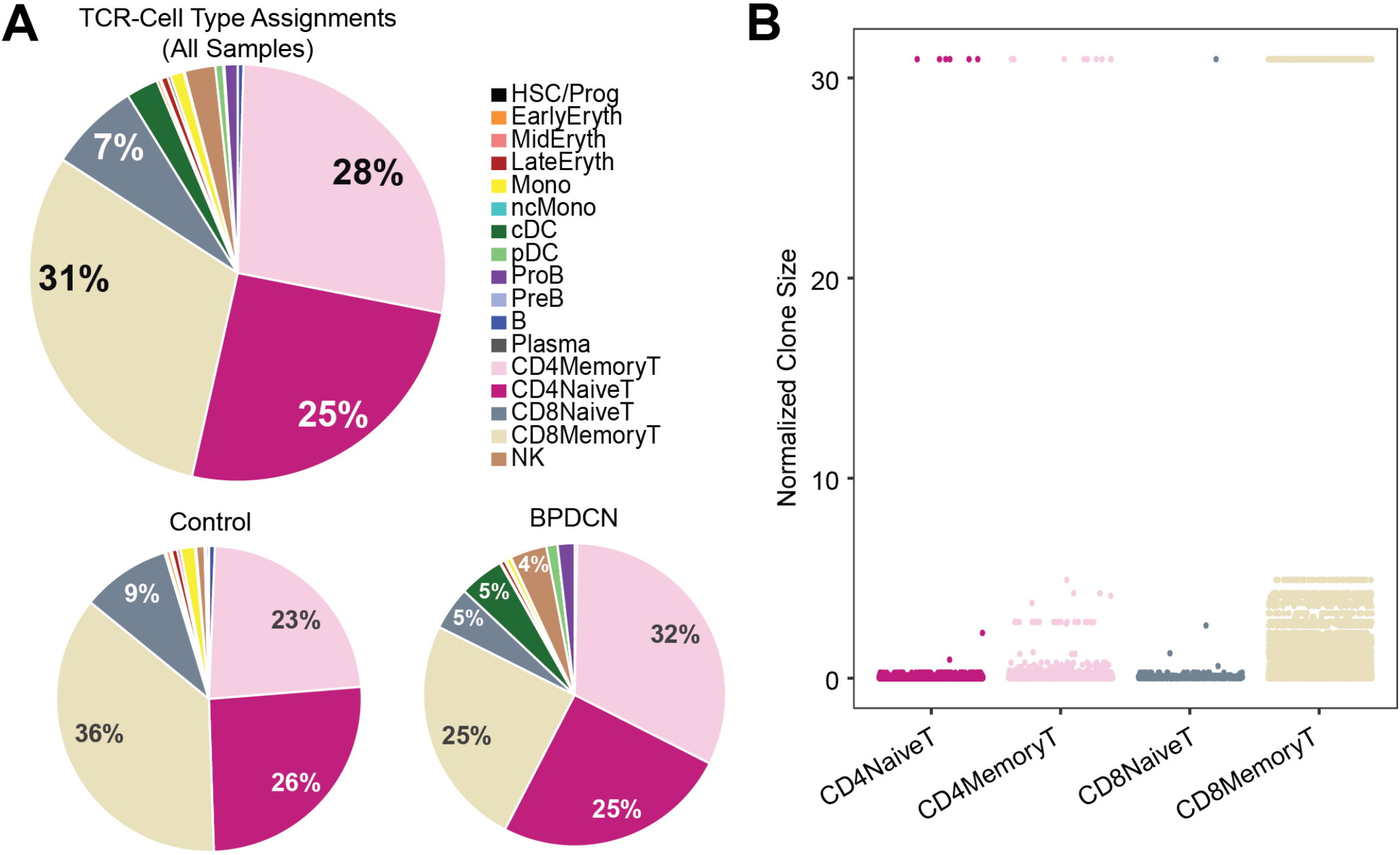
TCR sequences are detected in T-cells and clonally expanded in CD8+ Memory T-cells. (A) Pie charts show cell type assignments of cells in which a TCR sequence was detected. (B) Dot plot shows normalized clone size of all the T-cells in which a TCR sequence was detected. The distribution of values differs between T-cell subsets (*P* < 2.2E-16).

We next evaluated T-cell exhaustion by scoring each T-cell for the expression of exhaustion-associated genes (21). We overlaid cell type label, exhaustion scores, and normalized clone size parameters onto UMAP plots for each BPDCN sample containing T-cells (**Figure 7**). CD8+ Memory T-cell clusters were enriched for exhausted cells and expanded clonotypes in all patients, which was confirmed by quantification of signature scores and clone sizes in cell types (**Figure 8A**). Overall, exhaustion scores in CD8+ Memory T-cells in BPDCN 1-4 were significantly higher than controls (*P* = 0.0159), though the range of scores among individual cells within each sample was high (**Figure 8B**). These results are consistent with the pathway analysis above, as IFNA-mediated immune stimulation and low levels of TNFA signaling were previously associated with T-cell exhaustion (7,8,43).

**Figure 7:**
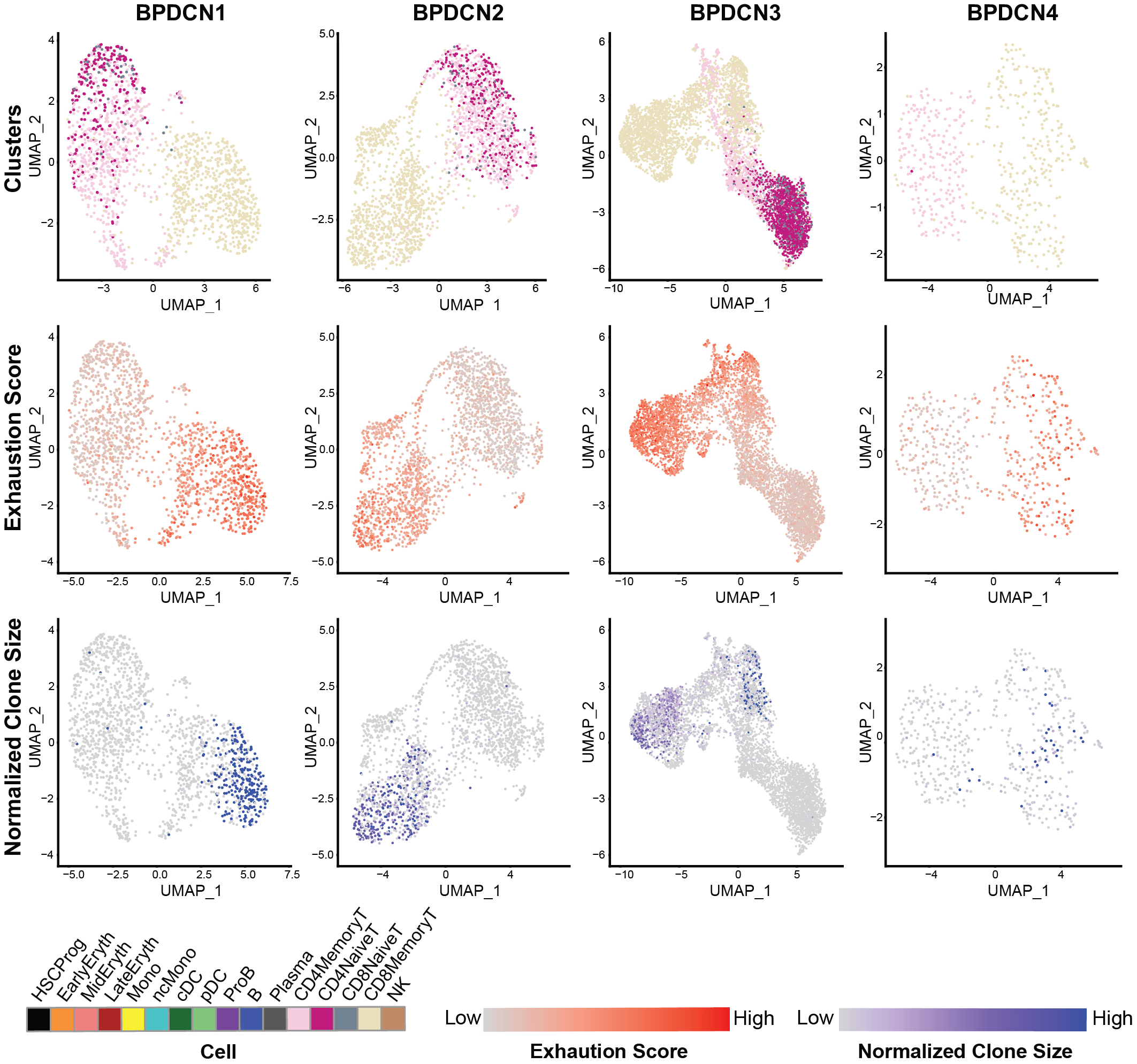
CD8+ Memory T-cells exhibit high expression of T-cell exhaustion-associated genes and larger TCR clones. UMAP plots of T-cell populations in each BPDCN sample that contains T-cells (BPDCN 1-4), separated by column. UMAPs are colored by cluster (first row); T-cell exhaustion score, with red indicating high exhaustion and gray indicating low exhaustion (second row); and normalized clone size, percentage of all cells in the dataset that share the same TCR sequence, with blue indicating high clone size and gray indicating low clone size (third row).

**Figure 8:**
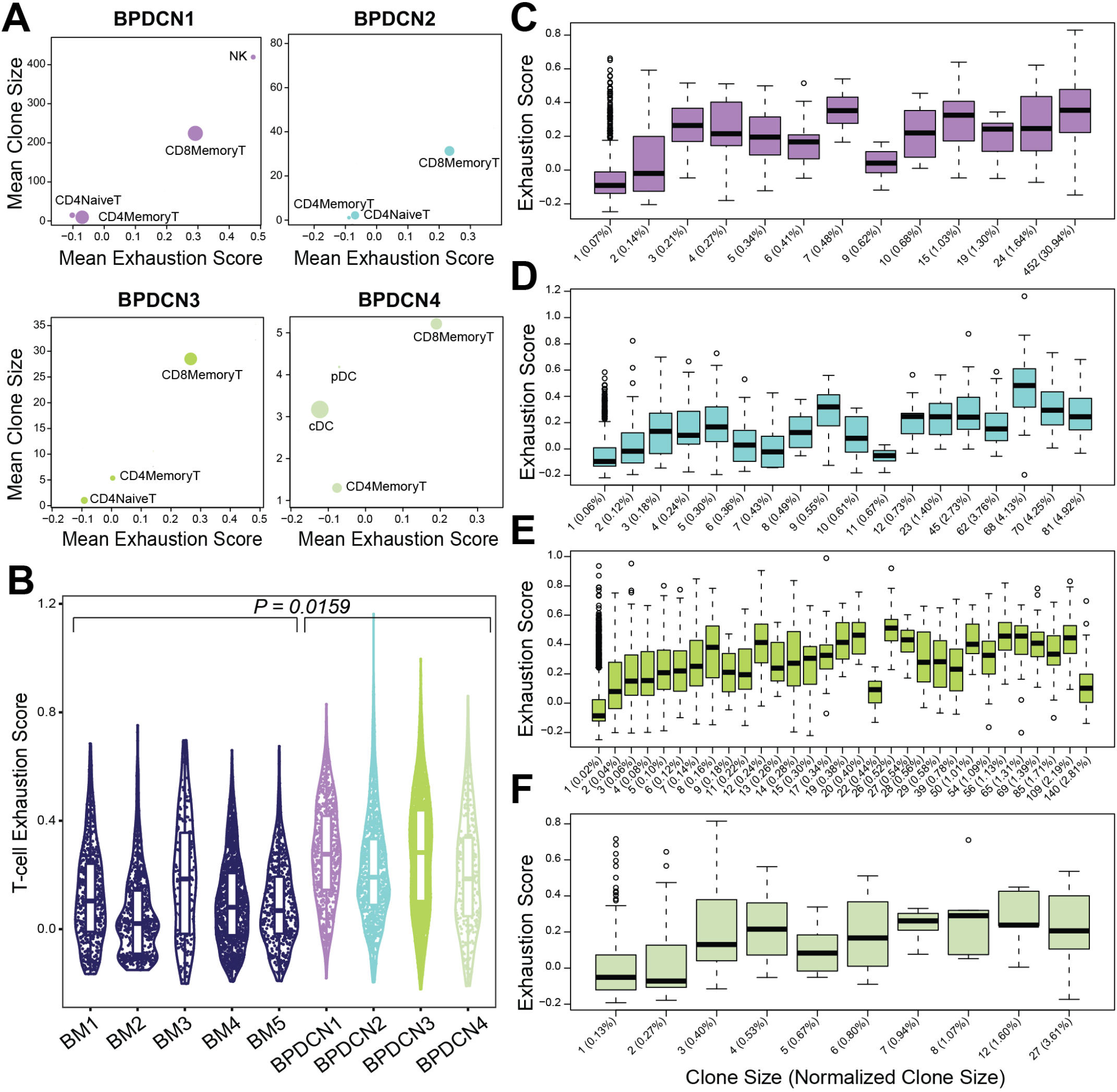
Exhaustion scores positively correlate with clone size in BPDCN samples. (A) Scatter plots show mean exhaustion score per cluster (x-axis) versus mean clone size (y-axis). Dots are scaled by the number of cells of each cluster in the dataset. (B) Violin plot shows T-cell exhaustion signature scores in CD8+ Memory T-cells in healthy controls (BM 1-5) and BPDCN samples with T-cells (BPDCN 1-4). (C-F) Box plots of cells binned by TCR clone size (x-axis) versus exhaustion score (y-axis) for T-cells in each BPDCN 1-4 sample as C-F. Parenthetical values on the x-axis labels indicate the normalized clone size.

To explore the relationship between exhaustion and clonotype sizes, we correlated these two parameters at the single-cell level. We found that CD8+ Memory T-cell clonotypes of various sizes can express T-cell exhaustion signature genes, including some clonotypes that make up a small proportion of the total T-cell population (**Figure 8C-F**). Spearman ranked correlation analysis revealed a significant positive relationship between clonotype size and exhaustion signature expression: rho of 0.67, 0.61, 0.63, and 0.4 for BPDCN 1-4, respectively (**Supplemental Figure 3**). These results suggest that clonal expansion and exhaustion of CD8+ Memory T-cell clonotypes may be functionally linked in the BPDCN tumor environment.

## 4 Discussion

To study the immune environment in BPDCN, we utilized paired single-cell sequencing of the transcriptome and T-cell receptors in healthy controls and patient samples. We identified 17 transcriptionally unique cell clusters in an integrated dataset of bone marrow from five younger and older healthy donors. We used this dataset to label cell types in five BPDCN patient bone marrow samples using four distinct classification algorithms, revealing heterogeneity in cell type proportions that correspond to clinical assessment of bone marrow involvement. An analysis of differentially regulated gene sets revealed that IFNA response genes were significantly increased in T-cells of the BPDCN samples and TNFA signaling genes were decreased. A deeper examination of expression within each gene set identified a subset of genes driving this enrichment, with the associated functions of T-cell proliferation, activation, and antigen presentation. TREK-seq showed T-cell specific TCR expression and uniformly distributed clone sizes in healthy controls; both of these results were more variable in BPDCN patient samples, suggesting possible TCR expression by tumor cells and antitumor reactivity by expanded T-cells in BPDCN. Further, we saw increased T-cell exhaustion in CD8+ Memory T-cells that positively correlated with increasing clone sizes. These results demonstrate the utility of single-cell multiomics by establishing a resource of the BPDCN tumor environment that provides new insights relevant to immuno-oncology.

Our results contribute to the body of research surrounding the role of the adaptive immune response in cancer and the relationship between T-cell receptor clonality and exhaustion. In a healthy immune system, activated pDCs produce large quantities of IFNA to stimulate activation of T-cells (10). In this study, we observe increased expression of IFNA response genes in T-cells relative to T-cells in the healthy controls, initially suggesting higher levels of IFNA production by pDC-like tumor cells. However, previous research on BPDCN has shown that pDC-like tumor cells produce lower levels of IFNA relative to pDCs from healthy individuals (44,45). Further, little research has been reported describing the role of TNFA signaling in BPDCN, though studies have shown increased TNFA signaling in the plasma of patients with acute myeloid leukemia (46,47). While the directionality of differential expression in these gene sets in T-cells is in contrast to what has been observed in the adjacent BPDCN cells, it is consistent with our findings of increased T-cell exhaustion. We hypothesize that, despite the tumor cells exhibiting lower IFNA production at the individual cell level, abnormal accumulation of pDC-like tumor cells in BPDCN may lead to increased IFNA production and chronic T-cell activation, eventually leading to T-cell exhaustion and consequent TNFA downregulation.

Exhaustion of cytotoxic T-cells has been a major hurdle in establishing effective immunotherapy treatments for blood cancers (24,25,48). Reactivation of T-cell effector function in exhausted T-cells is the goal of ICB, but aberrant IFNA and IFNG signaling have been shown to inhibit this process (49,50). While the prevalence of exhaustion has been studied in other cancers including acute myeloid leukemia, its role in BPDCN remains unknown. In this study, we establish that BPDCN patients have heightened T-cell exhaustion relative to controls. We also find a positive association between T-cell exhaustion and T-cell clonal expansion, consistent with other cancers (21,51). A lack of TCR expression by malignant cells is one feature that distinguishes BPDCN from mature T-cell malignancies, such as cutaneous T-cell lymphoma (52,53). However, little research has been done on T-cell biology in BPDCN, and none at the single-cell level. Our findings highlight a potential mechanism of T-cell exhaustion via persistent IFNA signaling that might be targeted to restore anti-tumor immunity.

Multimodal single-cell technologies enable investigation of gene regulation at a high resolution through multiple modalities representing interacting processes within a cell. This abundance of information allows us to deeply examine a small number of samples, however application of these methods to a larger patient population is prohibitively expensive and labor-intensive. For this study, we selected five BPDCN patients with varying levels of clinical bone marrow involvement, though more samples will be needed to ensure that the full spectrum of patient heterogeneity is captured. We observe interesting interactions between T-cell exhaustion and TCR expansion using parallel measurement of gene expression and TCR sequences, but other multimodal technologies could elucidate additional mechanisms of regulation in BPDCN. For example, the application of single-cell genotyping methods in future work would help to classify and characterize the transcriptomes of malignant tumor cells (2,16). Inclusion of additional patient samples and complementary single-cell measurements would strengthen our initial findings and uncover new results that further illuminate T-cell biology in BPDCN.

In summary, we apply scRNA-seq and TCR sequencing with computational techniques to catalog the cellular heterogeneity in BPDCN patient samples. Multiomics technology allows us to gain a deeper understanding of immune cell dynamics by assessing the diversity of immune cell states via scRNA-seq and the expansion of T-cell clonotypes through TREK-seq. Our results suggest that the balance between IFNA and TNFA signaling is disrupted in BPDCN, potentially leading to T-cell clonotype expansion and exhaustion. The discovery of mechanisms by which BPDCN cells evade immune destruction will lead to the development of new cancer therapies that leverage tumor-reactive T-cells.

## 6 Author Contributions

E.A.K.D., D.S., M.A., J.G., J.N., M.V., C.P.C., A.K.S. and P.v.G. conducted experiments and analyzed the data. E.A.K.D., A.A.L., and P.v.G designed the study and interpreted the data. G.K.G., A.A.L., H.R.M., and S.F.A. provided patient specimens and clinical perspectives. E.A.K.D. and P.v.G. wrote the manuscript. All authors edited the manuscript.

## 8 Acknowledgments

We thank patients for donating cells. We thank Phillip Dexheimer for helpful discussions and Patricia Rogers and the Broad Institute Flow Facility for technical support.

## 10 Data Availability Statement

Raw and processed data generated for this study will be available in the Gene Expression Omnibus (GEO) at https://www.ncbi.nlm.nih.gov/geo/. The scripts used to generate the results in this paper can be found at the project GitHub repository: https://github.com/EDePasquale/BPDCN.

