## Supplemental Figures and Tables for "Single-cell Multiomics Reveals Clonal T-cell Expansions and Exhaustion in Blastic Plasmacytoid Dendritic Cell Neoplasm"

### *Supplementary Material*

#### 1 Supplementary Figures

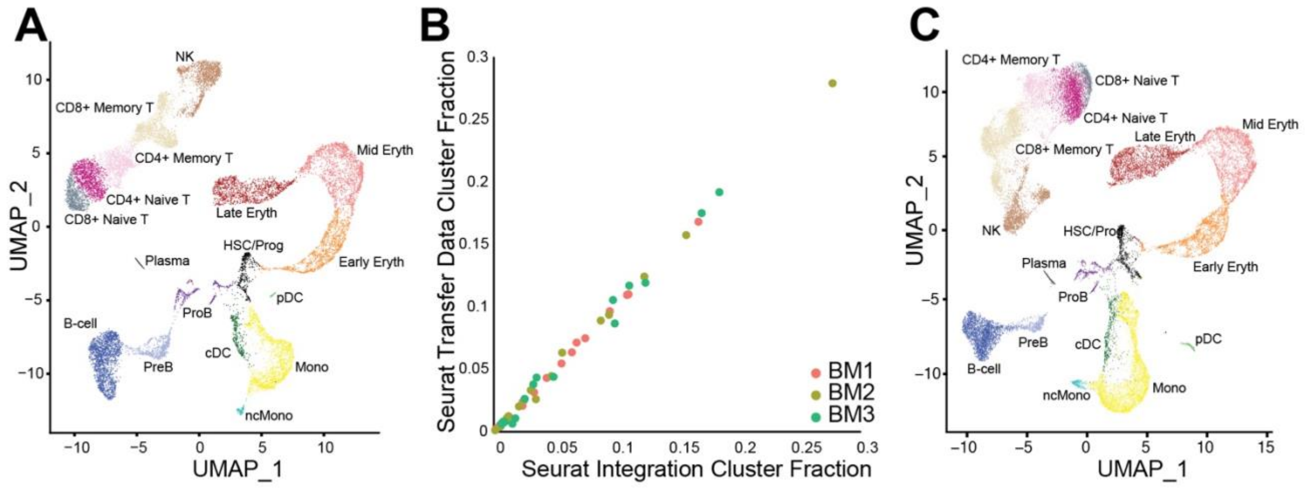

**Supplemental Figure 1: Integration and merging of healthy control samples.**

(A) UMAP visualization of Seurat integrated scRNA-seq data for hematopoietic cells from BM 1-3.
(B) Plot of the fraction of each cell type from each sample as classified by Seurat Integration (x-axis) and Seurat TransferData (y-axis). Colors represent samples, where each sample has up to 17 cell types present as dots on the graph.
(C) UMAP of Seurat clustering and sub-clustering of all BM samples (BM 1-5) identified 17 clusters of cells with similar transcriptional states following merging of BM 4-5 with the BM 1-3 reference from (A). Colors for each cell type in the UMAP plots represent cell types identified in the figure legend for **Figure 1**.

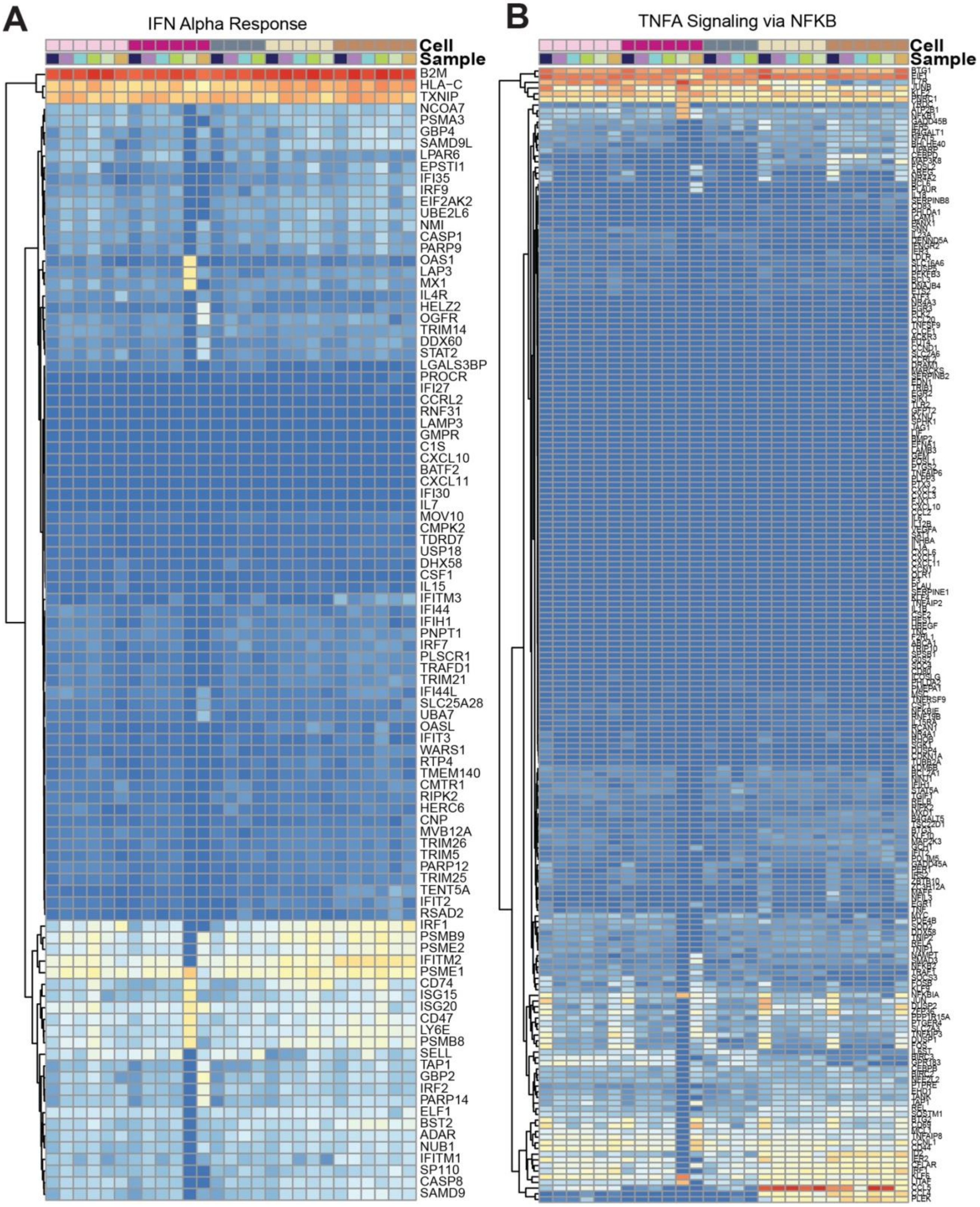

**Supplemental Figure 2: Expression of full IFNA Response and TNFA Signaling via NFKB gene sets in T and NK cells.**

(A) Heatmap shows log expression values for genes in the IFN Alpha signaling gene set (rows) for each sample and cell type (columns), clustered by row. Red indicates higher expression and blue indicates lower expression.
(B) Heatmap shows log expression values for genes in the TNFA Signaling via NFKB gene set. Colors for each cell type and sample match those in the figure legend for **Figure 4**.

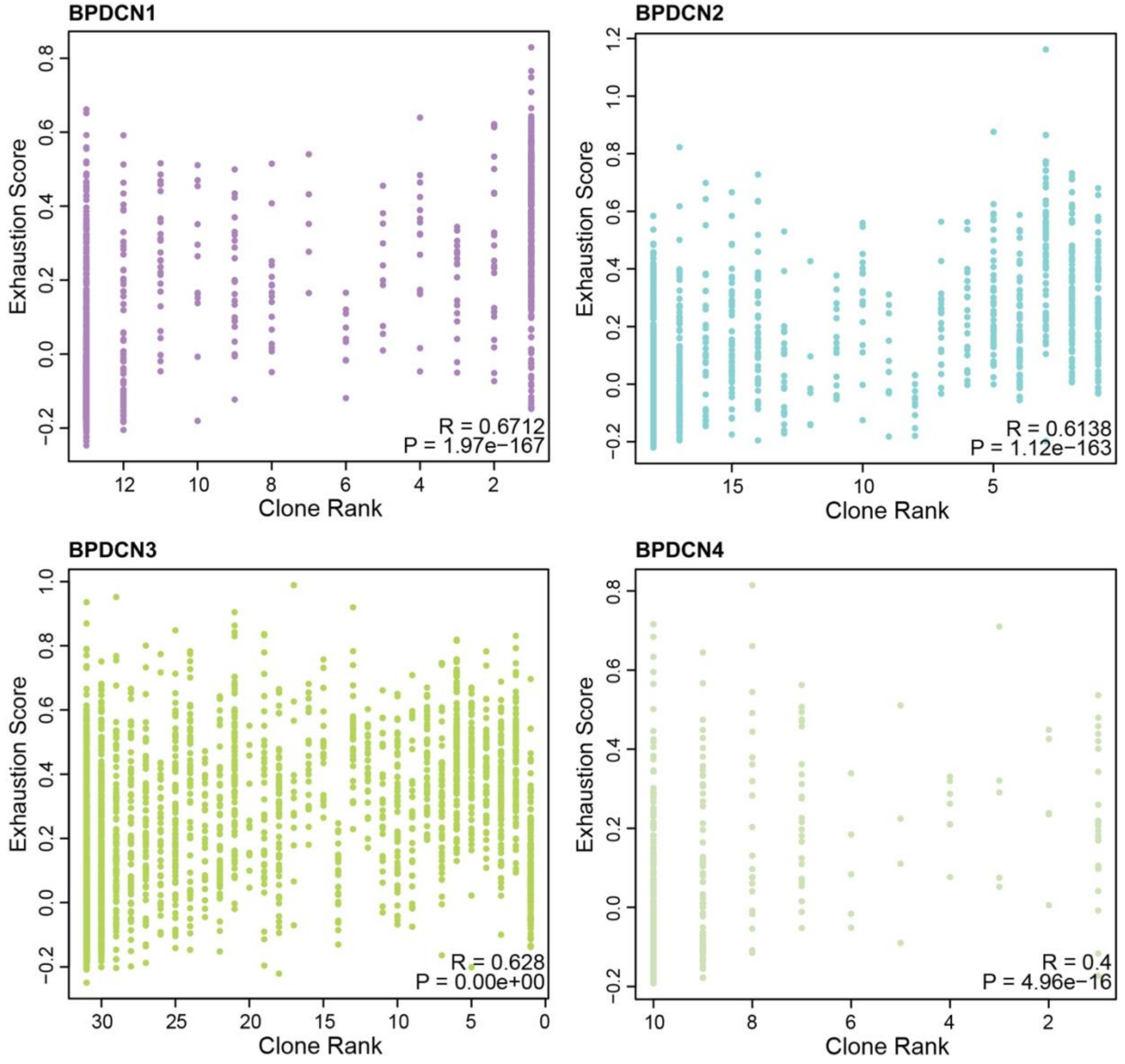

**Supplemental Figure 3: Clone size positively correlates with exhaustion score at the cell level in BPDCN T-cells.**

Correlation plots of T-cells for each of the BPDCN 1-4 samples, with each dot representing a cell. Clone rank (x-axis) of 1 means the most expanded clone of the dataset, with increasing rank following decreasing size. Exhaustion scores per cell are indicated by the y-axis. Spearman correlation values (R) and P-values (P) for the correlation are displayed in the bottom right corner of each plot.

#### 2 Supplementary Tables

**Supplemental Table 1: Donor and patient demographics.**

| Sample ID | Gender | Age | Tissue | Time | Bone marrow involvement | Cells After QC |
|---|---|---|---|---|---|---|
| BM1 | M | 31 | Bone marrow | — | — | 7635 |
| BM2 | F | 37 | Bone marrow | — | — | 2222 |
| BM3 | M | 45 | Bone marrow | — | — | 5219 |
| BM4 | M | 75 | Bone marrow | — | — | 7450 |
| BM5 | M | 74 | Bone marrow | — | — | 3200 |
| BPDCN1 | M | 70 | Bone marrow | Diagnosis | 30% on core biopsy, 1.5% by flow | 2869 |
| BPDCN2 | M | 63 | Bone marrow | Diagnosis | 10% | 4837 |
| BPDCN3 | M | 68 | Bone marrow | Diagnosis | 10%, based on aspirate/flow/RHP inference | 9818 |
| BPDCN4 | M | 64 | Bone marrow | Diagnosis | 97% | 4707 |
| BPDCN5 | M | 65 | Bone marrow | Diagnosis | 10-30% | 4846 |

**Supplemental Table 2: Primer sequences.**

| Name | Sequence (5' to 3') | Scale | Purification | Note |
|---|---|---|---|---|
| Human_TRBC | /5BiosG/GTGTTCCCACCCRAGGTCGCTGTGTTTGAGCCATCAGAAGCAGAGATCTCCCACACCCAAAAGGCCACACTGGTGTGCCTGGCCACAGGC | 4nmU | STD | Probe for enrichment; from Tu et al., 2019 |
| Human_TRAC | /5BiosG/CTGTCTGCCTATTCACCGATTTTGATTCTCAAACAAATGTGTCACAAAGTAAGGATTCTGATGTGTATATCACAGACAAAACTGTGCTAG | 4nmU | STD | Probe for enrichment; from Tu et al., 2019 |
| UPS2-N70x | CAAGCAGAAGACGGCATACGAGATGTCTCGTGGGCTCGG | 100nm | HPLC | Primer for TCR2 UPS2 amplification |
| 10X_SI-PCR_P5 | AATGATACGGCGACCACCGAGATCTACACTCTTTCCCTACACGACGCTC | 100nm | HPLC | Primer for TCR2 UPS2 amplification |
| aTCR-Seq | AGAGTCTCTCAGCTGGTACACGGCAGGGTCAGGITTCTGGATAT | 100nm | HPLC | Custom "index" sequencing primer for TCR-Seq |
| bTCR-Seq | CAAACACAGCGACCTCGGGTGGGAACACSTTKTTCAGGTCCT | 100nm | HPLC | Custom "index" sequencing primer for TCR-Seq |
| TRAVl | TCGTGGGCTCGGAGATGTGTATAAGAGACAGAGGTCGTTTTTCTTCATTCCTTAGTC | 25nm | STD | |
| TRAV2 | TCGTGGGCTCGGAGATGTGTATAAGAGACAGACGATACAACATGACCTATGAACGG | 25nm | STD | |
| TRAV3.1 | TCGTGGGCTCGGAGATGTGTATAAGAGACAGCTTTGAAGCTGAATTTAACAAGAGCC | 25nm | STD | |
| TRAV4.1 | TCGTGGGCTCGGAGATGTGTATAAGAGACAGCTCCCTGTTTATCCCTGCCGAC | 25nm | STD | |
| TRAV5.1 | TCGTGGGCTCGGAGATGTGTATAAGAGACAGAAACAAGACCAAAGACTCACTGTTC | 25nm | STD | |
| TRAV6 | TCGTGGGCTCGGAGATGTGTATAAGAGACAGAAGACTGAAGGTCACCTTTGATACC | 25nm | STD | |
| TRAV7 | TCGTGGGCTCGGAGATGTGTATAAGAGACAGACTAAATGCTACATTACTGAAGAATGG | 25nm | STD | |
| TRAV8 | TCGTGGGCTCGGAGATGTGTATAAGAGACAGGCATCAACGGTTTTGAGGCTGAATTTAA | 25nm | STD | |
| TRAV9 | TCGTGGGCTCGGAGATGTGTATAAGAGACAGGAAACCACTTCTTTCCACTTGGAGAA | 25nm | STD | |
| TRAV10 | TCGTGGGCTCGGAGATGTGTATAAGAGACAGTACAGCAACTCTGGATGCAGACAC | 25nm | STD | |
| TRAV12 | TCGTGGGCTCGGAGATGTGTATAAGAGACAGGAAGATGGAAGGTTTACAGCACA | 25nm | STD | |
| TRAV13.1 | TCGTGGGCTCGGAGATGTGTATAAGAGACAGGACATTCGTTCAAATGTGGGCGAA | 25nm | STD | |
| TRAV13.2 | TCGTGGGCTCGGAGATGTGTATAAGAGACAGGGCAAGGCCAAAGAGTCACCGT | 25nm | STD | |

| | | | |
|---|---|---|---|
| TRAV14 | TCGTGGGCTCGGAGATGTGTATAAGAGACAGTCCAGAAGGCAAGAAAATCCGCCA | 25nm | STD |
| TRAV16 | TCGTGGGCTCGGAGATGTGTATAAGAGACAGGCTGACCTTAACAAAGGCGAGACA | 25nm | STD |
| TRAV17 | TCGTGGGCTCGGAGATGTGTATAAGAGACAGTTAAGAGTCACGCTTGACACTTCCA | 25nm | STD |
| TRAV18 | TCGTGGGCTCGGAGATGTGTATAAGAGACAGGCAGAGGTTTTCAGGCCAGTCCT | 25nm | STD |
| TRAV19 | TCGTGGGCTCGGAGATGTGTATAAGAGACAGTCCACCAGTTCCTTCAACTTCACC | 25nm | STD |
| TRAV20 | TCGTGGGCTCGGAGATGTGTATAAGAGACAGGCCACATTAACAAAGAAGGAAAGCT | 25nm | STD |
| TRAV21 | TCGTGGGCTCGGAGATGTGTATAAGAGACAGGCCTCGCTGGATAAATCATCAGGA | 25nm | STD |
| TRAV22 | TCGTGGGCTCGGAGATGTGTATAAGAGACAGACGACTGTCGCTACGGAACGCTA | 25nm | STD |
| TRAV23 | TCGTGGGCTCGGAGATGTGTATAAGAGACAGCACAATCTCCTTCAATAAAAGTGCCA | 25nm | STD |
| TRAV24 | TCGTGGGCTCGGAGATGTGTATAAGAGACAGACGAATAAGTGCCACTCTTAATACCA | 25nm | STD |
| TRAV25 | TCGTGGGCTCGGAGATGTGTATAAGAGACAGGTTTGGAGAAGCAAAAAAGAACAGCT | 25nm | STD |
| TRAV26.1 | TCGTGGGCTCGGAGATGTGTATAAGAGACAGCAGAAGACAGAAAGTCCAGCACCT | 25nm | STD |
| TRAV26.2 | TCGTGGGCTCGGAGATGTGTATAAGAGACAGATCGCTGAAGACAGAAAGTCCAGT | 25nm | STD |
| TRAV27 | TCGTGGGCTCGGAGATGTGTATAAGAGACAGACTAACCTTTCAGTTTGGTGATGCAA | 25nm | STD |
| TRAV29 | TCGTGGGCTCGGAGATGTGTATAAGAGACAGCTTAAACAAAAGTGCCAAGCACCTC | 25nm | STD |
| TRAV30 | TCGTGGGCTCGGAGATGTGTATAAGAGACAGAATATCTGCTTCATTTAATGAAAAAAAGC | 25nm | STD |
| TRAV34 | TCGTGGGCTCGGAGATGTGTATAAGAGACAGCCAAGTTGGATGAGAAAAAGCAGCA | 25nm | STD |
| TRAV35 | TCGTGGGCTCGGAGATGTGTATAAGAGACAGCTCAGTTTGGTATAACCAGAAAGGA | 25nm | STD |
| TRAV36 | TCGTGGGCTCGGAGATGTGTATAAGAGACAGGGAAGACTAAGTAGCATATTAGATAAG | 25nm | STD |
| TRAV38 | TCGTGGGCTCGGAGATGTGTATAAGAGACAGCTGTGAACTTCCAGAAAGCAGCCA | 25nm | STD |
| TRAV39 | TCGTGGGCTCGGAGATGTGTATAAGAGACAGCCTCACTTGATACCAAAGCCCGT | 25nm | STD |
| TRAV40 | TCGTGGGCTCGGAGATGTGTATAAGAGACAGAGGCGGAAATATTAAAGACAAAAACTC | 25nm | STD |
| TRAV41 | TCGTGGGCTCGGAGATGTGTATAAGAGACAGGATTAATTGCCACAATAAACATACAGG | 25nm | STD |

| | | | |
|---|---|---|---|
| TRBV2 | TCGTGGGCTCGGAGATGTGTATAAGAGACAGGCCTGATGGATCAAATTTCACTCTG | 25nm | STD |
| TRBV3-1 | TCGTGGGCTCGGAGATGTGTATAAGAGACAGTCTCACCTAAATCTCCAGACAAAGCT | 25nm | STD |
| TRBV4 | TCGTGGGCTCGGAGATGTGTATAAGAGACAGCCTGAATGCCCCAACAGCTCTC | 25nm | STD |
| TRBVS-4,8 | TCGTGGGCTCGGAGATGTGTATAAGAGACAGCTCTGAGCTGAATGTGAACGCCT | 25nm | STD |
| TRBVS-1 | TCGTGGGCTCGGAGATGTGTATAAGAGACAGCGATTCTCAGGGCGCCAGTTCTCT | 25nm | STD |
| TRBV6-1 | TCGTGGGCTCGGAGATGTGTATAAGAGACAGTGGCTACAATGTCTCCAGATTAAACAA | 25nm | STD |
| TRBV6-2,3 | TCGTGGGCTCGGAGATGTGTATAAGAGACAGCCCTGATGGCTACAATGTCTCCAGA | 25nm | STD |
| TRBV6-4 | TCGTGGGCTCGGAGATGTGTATAAGAGACAGGTGTCTCCAGAGCAAACACAGATGATT | 25nm | STD |
| TRBV6-5,6 | TCGTGGGCTCGGAGATGTGTATAAGAGACAGGTCTCCAGATCAACCACAGAGGAT | 25nm | STD |
| TRBV6-8 | TCGTGGGCTCGGAGATGTGTATAAGAGACAGGTCTCTAGATTAAACACAGAGGATTTC | 25nm | STD |
| TRBV6-9 | TCGTGGGCTCGGAGATGTGTATAAGAGACAGGGCTACAATGTATCCAGATCAAACA | 25nm | STD |
| TRBV7-2 | TCGTGGGCTCGGAGATGTGTATAAGAGACAGTCGCTTCTCTGCAGAGAGGACTGG | 25nm | STD |
| TRBV7-3 | TCGTGGGCTCGGAGATGTGTATAAGAGACAGCGGTTCTTTGCAGTCAGGCCTGA | 25nm | STD |
| TRBV7-8 | TCGTGGGCTCGGAGATGTGTATAAGAGACAGCCAGTGATCGCTTCTTTGCAGAAA | 25nm | STD |
| TRBV?-4,6 | TCGTGGGCTCGGAGATGTGTATAAGAGACAGTCTCCACTCTGAMGATCCAGCGCA | 25nm | STD |
| TRBV7-7 | TCGTGGGCTCGGAGATGTGTATAAGAGACAGGCAGAGAGGCCTGAGGGATCCAT | 25nm | STD |
| TRBV7-9 | TCGTGGGCTCGGAGATGTGTATAAGAGACAGCTGCAGAGAGGCCTAAGGGATCT | 25nm | STD |
| TRBV9 | TCGTGGGCTCGGAGATGTGTATAAGAGACAGCTCCGCACAACAGTTCCCTGACTT | 25nm | STD |
| TRBV10-1,3 | TCGTGGGCTCGGAGATGTGTATAAGAGACAGCAGATGGCTAYAGTGTCTCTAGATCAAA | 25nm | STD |
| TRBV10-2 | TCGTGGGCTCGGAGATGTGTATAAGAGACAGGTTGTCTCCAGATCCAAGACAGAGAA | 25nm | STD |
| TRBV11 | TCGTGGGCTCGGAGATGTGTATAAGAGACAGGCAGAGAGGCTCAAAGGAGTAGACT | 25nm | STD |
| TRBV12-3,4 | TCGTGGGCTCGGAGATGTGTATAAGAGACAGGCTAAGATGCCTAATGCATCATTCTC | 25nm | STD |
| TRBV12-5 | TCGTGGGCTCGGAGATGTGTATAAGAGACAGCTCAGCAGAGATGCCTGATGCAACT | 25nm | STD |

| | | | |
|---|---|---|---|
| TRBV13 | TCGTGGGCTCGGAGATGTGTATAAGAGACAGTCTCAGCTCAACAGTTCAGTGACTA | 25nm | STD |
| TRBV14 | TCGTGGGCTCGGAGATGTGTATAAGAGACAGGCTGAAAGGACTGGAGGGACGTAT | 25nm | STD |
| TRBV15 | TCGTGGGCTCGGAGATGTGTATAAGAGACAGGATAACTTCCAATCCAGGAGGCCG | 25nm | STD |
| TRBV16 | TCGTGGGCTCGGAGATGTGTATAAGAGACAGGCTAAGTGCCTCCCAAATTCACCC | 25nm | STD |
| TRBV18 | TCGTGGGCTCGGAGATGTGTATAAGAGACAGGGAACGATTTTCTGCTGAATTTCCCA | 25nm | STD |
| TRBV19 | TCGTGGGCTCGGAGATGTGTATAAGAGACAGGGTACAGCGTCTCTCGGGAGAAGA | 25nm | STD |
| TRBV20-1 | TCGTGGGCTCGGAGATGTGTATAAGAGACAGGGACAAGTTTCTCATCAACCATGCAA | 25nm | STD |
| TRBV24-1 | TCGTGGGCTCGGAGATGTGTATAAGAGACAGTGGATACAGTGTCTCTCGACAGGC | 25nm | STD |
| TRBV25-1 | TCGTGGGCTCGGAGATGTGTATAAGAGACAGCAACAGTCTCCAGAATAAGGACGGA | 25nm | STD |
| TRBV27-1 | TCGTGGGCTCGGAGATGTGTATAAGAGACAGTACAAAGTCTCTCGAAAAGAGAAGAGGA | 25nm | STD |
| TRBV28 | TCGTGGGCTCGGAGATGTGTATAAGAGACAGGGGGTACAGTGTCTCTAGAGAGA | 25nm | STD |
| TRBV29 | TCGTGGGCTCGGAGATGTGTATAAGAGACAGGTTTCCCATCAGCCGCCCAAACCTA | 25nm | STD |
| TRBV30 | TCGTGGGCTCGGAGATGTGTATAAGAGACAGCAGACCCCAGGACCGGCAGTTCAT | 25nm | STD |
